# A widespread group of large plasmids in methanotrophic *Methanoperedens* archaea

**DOI:** 10.1101/2022.02.01.478723

**Authors:** Marie C. Schoelmerich, Heleen T. Oubouter, Rohan Sachdeva, Petar Penev, Yuki Amano, Jacob West-Roberts, Cornelia U. Welte, Jillian F. Banfield

## Abstract

Anaerobic methanotrophic (ANME) archaea conserve energy from the breakdown of methane, an important driver of global warming, yet the extrachromosomal genetic elements that impact the activities of ANME archaea are little understood. Here we describe large plasmids associated with ANME archaea of the *Methanoperedens* genus. These have been maintained in two bioreactors that contain enrichment cultures dominated by different *Methanoperedens* species and co-occur with *Methanoperedens* species in other anoxic environments. By manual curation we show that two of the plasmids are large (155,607 bp and 191,912 bp), circular, and replicate bidirectionally. The group of *Methanoperedens* species that carry these plasmids is related to “*Ca*. Methanoperedens nitroreducens”, “*Ca*. Methanoperedens ferrireducens”, “*Ca*. Methanoperedens manganicus" and the plasmids occur in the same copy number as the main chromosome. The larger plasmid encodes transporters that potentially enhance access to Ni, which is required for the methyl-CoM reductase (Mcr), Co required for the cobalamin cofactor needed for methyltransferases, and amino acid uptake. We show that many plasmid genes are actively transcribed, including genes involved in plasmid chromosome maintenance and segregation, a Co^2+^/Ni^2+^ transporter and cell protective proteins. Notably, one plasmid carries three tRNAs and two colocalized genes encoding ribosomal protein uL16 and elongation factor eEF2. These are not encoded in the host *Methanoperedens* genome and uL16 and eEF2 were highly expressed, indicating an obligate interdependence between this plasmid and its host. The finding of plasmids of *Methanoperedens* opens the way for the development of genetic vectors that could be used to probe little understood aspects of *Methanoperedens* physiology. Ultimately, this may provide a route to introduce or alter genes that may enhance growth and overall metabolism to accelerate methane oxidation rates.

## Introduction

Anaerobic oxidation of methane (AOM) is a microbial process of a polyphyletic group of archaea termed ANME. While most known ANME inhabit marine environments and rely on a syntrophic partner (ANME-1, ANME-2a-c, ANME-3), the *Methanoperedenaceae* (formerly ANME-2d) live in freshwater ecosystems and use nitrate, iron oxide, or manganese oxide as extracellular electron acceptors [1–4]. AOM has sparked increasing interest due to its role in naturally decreasing CH_4_ emissions by reoxidizing it to CO_2_. Methanogenic archaea (methanogens) make CH_4_ using either CO_2_, methylated compounds or acetate as the carbon source [5]. ANME seem to reverse the methanogenesis process by using largely the same enzymatic machinery in a process termed “reverse methanogenesis” [6,7]. Understanding their metabolism and how it is regulated is of increasing interest, due to their ecological importance in the global CH_4_ cycle.

The discovery of extrachromosomal elements (ECEs) in the archaeal domain of life is still in its infancy. 307 plasmid sequences from archaea make up less than 1% of all 34,908 plasmid sequences on NCBI (https://www.ncbi.nlm.nih.gov/genome/browse#!/plasmids/, Jan-18, 2022). Most plasmid sequences originate from halophilic archaea, which have been model organisms to study these archaeal ECEs [8,9]. Yet, there have also been several plasmids isolated and decrypted from a few methanogens [10,11]. Although the native function of these plasmids is not well established, they have been instrumental in paving the way towards genetically engineering methanogens.

The recent discovery of Borgs associated with methane-oxidizing members of the *Methanoperedens* has ignited interest in finding ways to understand and potentially leverage these novel ECEs for genetic engineering purposes [12]. Here, we describe the discovery of *Methanoperedens* plasmids in metagenomic datasets originating from two bioreactors as well as several natural ecosystems. We manually curated two plasmid genomes to completion. The genetic repertoire and expression profile of the plasmids is presented, and elements for a shuttle vector for future genetic engineering approaches are identified. We anticipate that this discovery will lead to important advances in understanding the ecology, physiology, biochemistry and bioenergetics of ANME archaea.

## Results

### Search for ECEs revealed large plasmids

To find plasmids that associate with *Methanoperedens* we searched for contigs with plasmid-like gene content and taxonomic profiles most similar to those of *Methanoperedens* but that were not part of a *Methanoperedens* chromosome in metagenomic datasets from two bioreactors that are dominated by “*Candidatus* Methanoperedens BLZ2” (Bioreactor 1, [13]) and “*Candidatus* Methanoperedens nitroreducens Vercelli” (Bioreactor 2, [14]). The bioreactors have been maintained since 2015 and the main metabolism of both enrichment cultures is nitrate-dependent AOM. Samples for DNA and RNA extractions were taken in April 2021 and again in October 2021. “*Ca.* Methanoperedens BLZ2” comprised ~44% of the sampled community in Bioreactor 1. It has a ~3.93 Mbp genome and coexists with *Methylomirabilis oxyfera* with a ~2.73 Mbp genome that accounted for 26% of the organisms in the sampled community, whereas all other organisms were <~5%. “*Ca.* Methanoperedens nitroreducens Vercelli” in Bioreactor 2 constituted ~78% of the sampled community. It has a ~3.28 MBp genome and coexists with many other microorganisms, each of which comprised <4% of the community.

We found two plasmids in Bioreactor 1: HMp_v1 and HMp_v5 and two plasmids in Bioreactor 2: HMp_v2 and HMp_v3 (**Table 1**), both of which are distinct from the plasmids in Bioreactor 1. Importantly, *Methanoperedens* are the only archaea that coexist with these archaeal plasmids and this enabled us to confidently assign “*Ca.* Methanoperedens BLZ2” (4357x coverage) as the host of HMp_v5 (4599x coverage) and “*Ca.* Methanoperedens nitroreducens Vercelli” (4204x coverage) as the host of HMp_v2 (5405x coverage). HMp_v3 (19x coverage) may be a plasmid of Mp_Bioreactor_2_Methanoperedens_40_26 (26x coverage) or a rare plasmid of the Vercelli strain. No alternative potential host was identified for HMp_v1 (27x coverage), so this may be a second rare plasmid of “*Ca.* Methanoperedens BLZ2”. Overall, we infer that the abundant plasmids are maintained at the same copy number as the *Methanoperedens* chromosome. This parallels findings for *Halobacteriales,* which usually have the same copy number of chromosomes and megaplasmids [15].

**Table 1.** Features of *Methanoperedens* plasmids.

| Plasmid | Ecosystem | Genome | Site | Size (bp) | % GC | Cov | # Ctg | # Features |
| --- | --- | --- | --- | --- | --- | --- | --- | --- |
| **v1** | Bioreactor | complete | Bioreactor 1 | 191,912 | 39.2 | 27 | 1 | 186 |
| **v2** | Bioreactor | complete | Bioreactor 2 | 155,607 | 39.2 | 5405 | 1 | 158 |
| **v3** | Bioreactor | partial | Bioreactor 2 | 63,092 | 38.9 | 19 | 1 | 56 |
| **v5** | Bioreactor | almost complete | Bioreactor 1 | 185,698 | 39.7 | 4599 | 1 | 164 |
| **v4** | Environmental | fragment | SRVP | 35,271 | 40.9 | 8 | 8 | 35 |
| **v6** | Environmental | fragment | Japan | 24,153 | 40.4 | 15 | 1 | 25 |
| **v7** | Environmental | partial | Rifle | 163,653 | 41.2 | 17 | 3 | 125 |
| **v8** | Environmental | almost complete | SRVP | 253,501 | 41.5 | 20 | 9 | 288 |

We then searched for additional sequences in our metagenomic database and identified four related plasmids from three different ecosystems (**Table 1, Figure 1A** and **Figure S1**). These sequences originated from the sedimentary rock Horonobe Japan Deep Subsurface research site [16], a shallow aquifer adjacent to the Colorado River (Rifle, CO, USA; [17]), and saturated wetland soil (SR, CA, USA; [18]). Thus, we suggest that plasmids may often be associated with certain *Methanoperedens* species. The Horonobe plasmid, HMp_v6, only co-occurs with one *Methanoperedens* species (Ig18389_08E140C01_z1_2020_Methanoperedens_40_15) that is at very similar coverage to the plasmids (both 15x coverage). The Rifle plasmid HMP_v7 (17x coverage) occurs in a sample with many archaea, but we only identified one as a *Methanoperedens* species, RBG_16_Methanoperedens_41_19 (19x coverage [17]). Thus, we also suspect a plasmid-host ratio of ~1:1 for these environmental plasmids.

**Figure 1.**
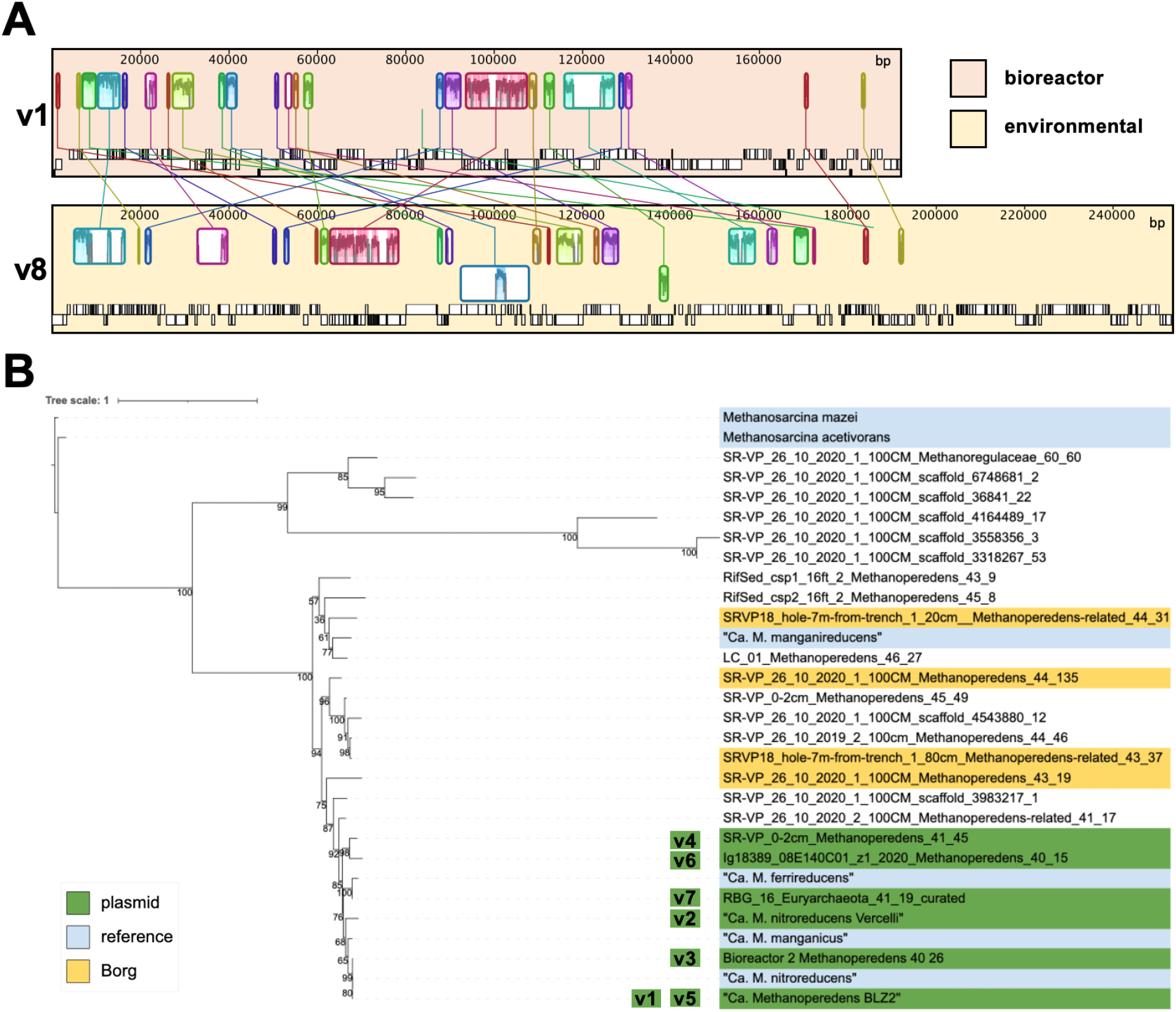
Nucleotide alignment of plasmid versions v1 and v8 and phylogenetic tree of different *Methanoperedens* species. **A.** All 9 contigs of HMp_v8 were sorted and aligned onto the complete genome of HMp_v1 using the MCM algorithm in Geneious. **B.** RpS3-based phylogenetic tree of *Methanoperedens* species rooted on RpS3 of the methanogen *M. mazei*. Plasmid host associations are inferred for plasmids 1-7 based on co-occurrence and same coverage.

We constructed a phylogenetic tree to examine the pattern of associations between various *Methanoperedens* species and plasmids. The bioreactors do not contain Borgs and the *Methanoperedens* in the bioreactors are not closely related to species that host Borgs (**Figure 1B**). Only in the case of HMp_v4 and HMp_v8 do Borgs and plasmids co-occur in samples with *Methanoperedens*, but these samples contain many *Methanoperedens* species. Notably, we find that the species of *Methanoperedens* that host the plasmids are phylogenetically clustered together and are distinct from the species suggested to host Borgs. Thus, this clade of *Methanoperedens* plausibly consistently hosts the plasmids. This *Methanoperedens* species group includes “*Ca.* M. nitroreducens”, “*Ca.* M. ferrireducens" and “*Ca.* M. manganicus”, so plasmids may also occur in the enrichment cultures that contain these strains.

### Curation and completion of two plasmid sequences

Two plasmid genomes were curated to completion (see Methods). After curation, the ends of each plasmid sequence were identical and spanned by paired reads, revealing that they are circular. The plasmids carry genes on both strands and most genes are within polycistronic transcription units. HMp_v1 is 155,607 bp and has 158 ORFs, HMp_v2 is 191,912 bp and has 178 ORFs. These two plasmids do not encode tRNAs, rRNAs or ribosomal proteins. Large stretches of v1 and v2 align, resulting in 139 shared (and mostly identical) proteins. Forty-seven proteins are unique to v1 and 19 proteins are unique to v2 (**Figure 2, Table S1**).

**Figure 2.**
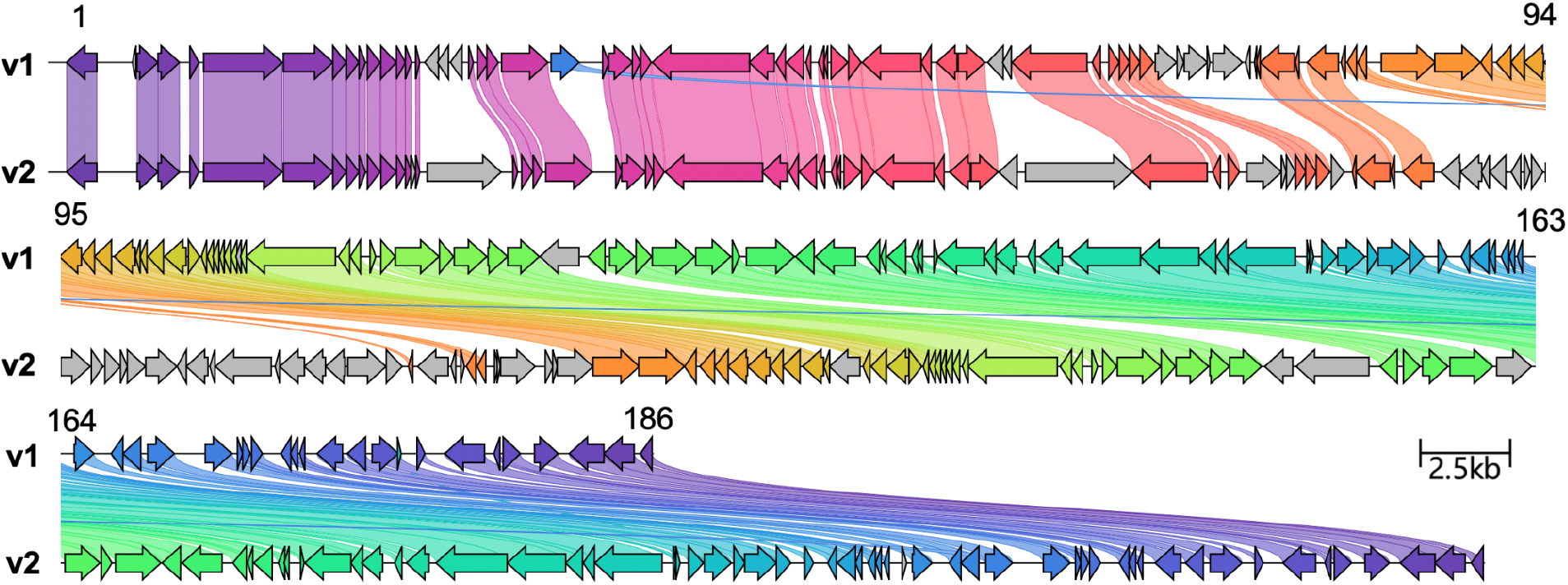
Alignment of curated HMp_v1 from Bioreactor 1 and HMp_v2 from Bioreactor 2. Colored genes highlight homologues based on amino acid identity. Gene numbers of HMp_v1 are indicated on top.

The partially curated HMp_v5 plasmid is encoded on a single 185,698 bp contig with 166 ORFs. It only aligns with HMp_v1 and HMp_v2 in some regions, indicating that it is more distantly related than v1 is to v2 (**Figure 3**). It encodes ribosomal protein uL16 (Bioreactor_1_104068_82). No uL16 gene was identified in *Methanoperedens* in the bioreactor or on any unbinned contigs in the metagenome. Encoded adjacent to uL16 on HMp_v5 is translation initiation factor 2 subunit beta (aeIF-2b) that also appears to be missing from the host genome. HMp_v5 also encodes tRNA Asp, tRNA Arg and tRNA Val. Interestingly, the host appears to lack tRNA Asp and the anticodons of the plasmid tRNA Val and tRNA Arg are not represented in the tRNA inventory of the host. The plasmid tRNAs group phylogenetically with tRNAs from other species of the same class as *Methanoperedens* (*Methanomicrobia*). Thus, the plasmid tRNAs likely derived from *Methanomicrobia* (**Figure S2-S4**).

**Figure 3.**
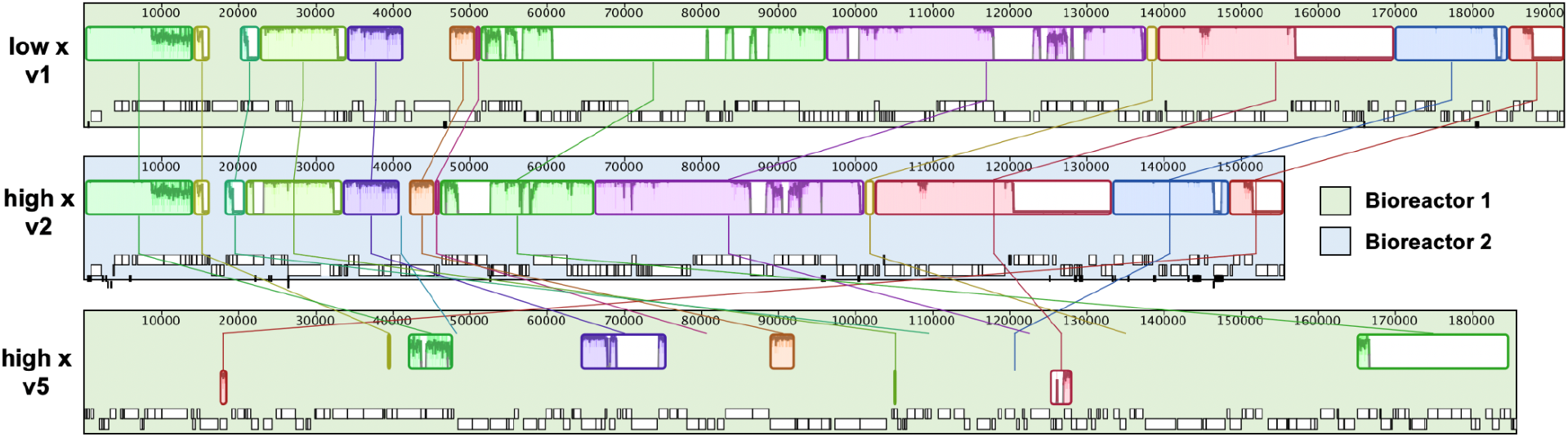
Genome alignment of curated plasmid versions v1, v2 and contig of v5. Genomes were aligned with the progressiveMauve algorithm in Geneious.

### Plasmid replication, stability and segregation

The two complete plasmid genomes had cumulative GC skew indicative of bi-directional replication (**Figure S5**). The origin of replication encodes an ATPase and the origin of replication recognition protein (Orc1/Cdc6) (**Figure 4A**). A Cdc24-bearing protein encoded elsewhere in the genome (ORF94; all orf numbers apply to the HMp_v1 version, unless indicated otherwise) may also be involved in replication initiation. ORF2 and ORF3 fall within protein subfamilies that include sequences loosely annotated as RepA. Modeling supports their annotation as RepA1 and RepA2 with closest structural similarity to two subunits of the trimerization core of human RepA (PDB: 1l1o:F and 1l1o:B) proteins, which are ssDNA binding proteins essential for preventing reannealing and degradation of the growing ssDNA chain during replication. The region encompassing the origin of replication and the adjacent genes encoding replication-associated proteins are likely important core elements if the plasmids are adapted into a genetic engineering vector.

**Figure 4.**
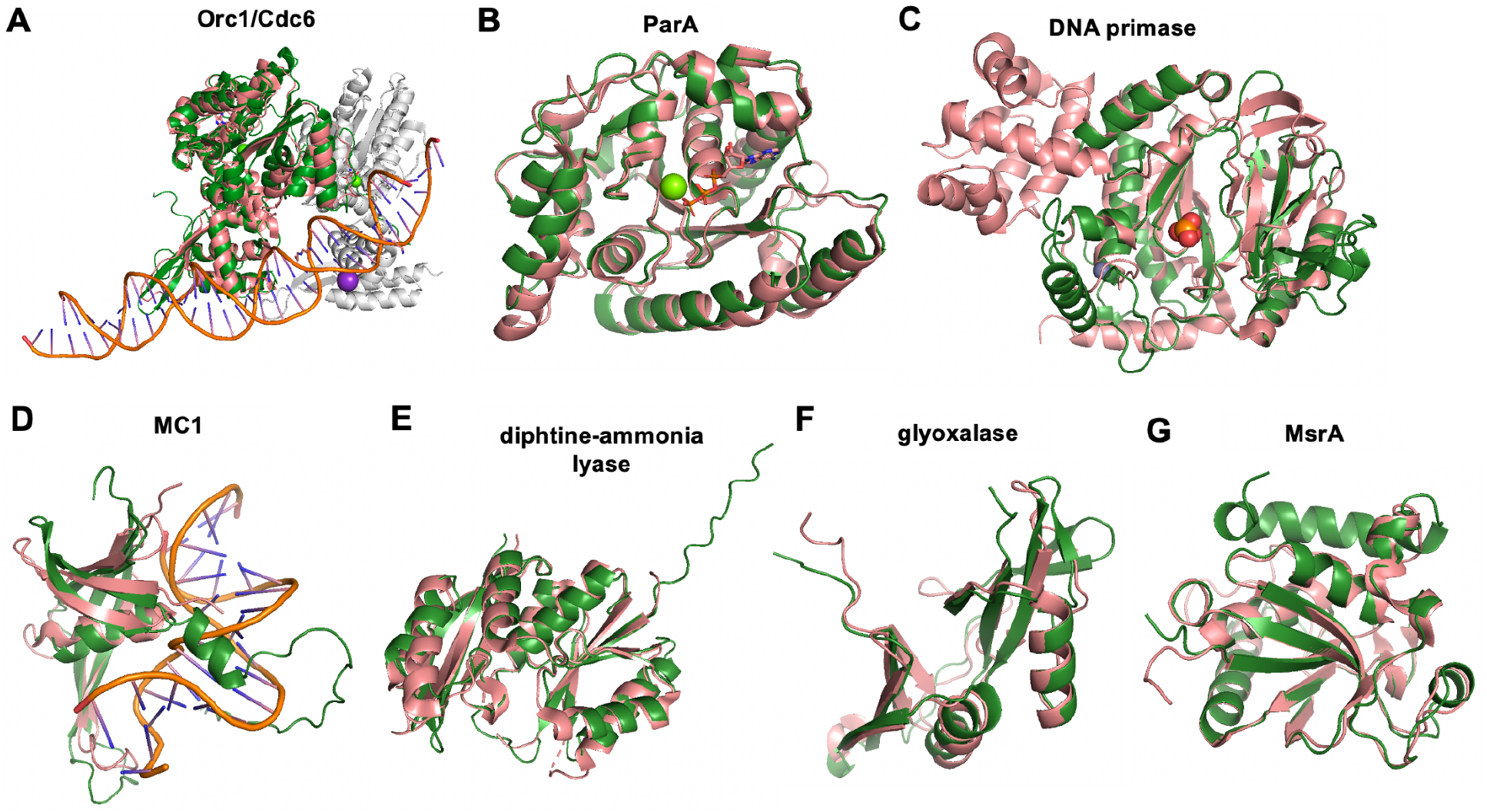
Predicted structures of plasmid proteins from HMp_v1. Plasmid proteins (green) were superimposed on best hits (salmon) from PDBeFold in pyMOL. **A.** Orc1/Cdc6 (ORF1) and heterodimeric Orc1-1/Orc1-3 complex from *Sulfolobus solfataricus* (PDB: 2QBY_A, RMSDpymol = 1.91). Orc1-3 (2QBY_B) is drawn in grey. **B.** ParA (ORF167) and HpSoj from *Helicobacter pylori* (6iuc_A, RMSD = 0.9). **C.** DNA primase (ORF120) and *P. furiosus* homologue (PDB: 1v33A, RMSD = 1.91). **D.** Nucleoid protein MC1 (ORF157) and DNA-archaeal MC1 protein complex from *M. thermophila* (2khl_A, RMSD = 2.30). **E.** DAL (ORF67) and *Pyrococcus horikoshii* homologue (PDB: 2d13_A, RMSD = 1.37). **F.** Glyoxalase (ORF36) and *Enterococcus faecalis* homologue (PDB: 2P25_A, RMSD = 1.19). **G.** MsrA (ORF81) and *Mycobacterium tuberculosis* homologue (PDB: 1nwa_A, RMSD = 0.75).

The plasmids encode six helicases with variable domain topologies. ORF5 encodes a helicase with an additional N-terminal N-6 methylase domain that may unwind DNA and immediately methylate nascent DNA at the replication fork. A ubiquitin-activating (adenylating) ThiF family protein and a RadC-like protein tied to recombinational repair at the replication fork [19] as well as a DUF488-bearing protein, nucleotide-sensing YpsA protein and UvrD helicase resembling Dna2 are also part of this genetic neighbourhood. RadC (ORF11) and the UvrD helicase (ORF20) occur in many of the plasmids and were thus used to phylogenetically determine their relatedness (**Figure S6A, B**). Indeed, the plasmid proteins clustered together, corroborating whole genome alignments that indicate they are related. Supplementing the sequences with top hits from BlastP search on NCBI furthermore substantiated that the plasmid proteins are most closely related to *Methanoperedens*.

Other plasmid genes are predicted to be involved in segregation of replicated genetic material. A structure-based homology search identified ParA (**Figure 4B**), but there is no obvious ParB or AspA homologue, which are the other two components of the tripartite DNA partitioning system in Crenarchaeota [20]. In both plasmids ParA is accompanied by a transposase and a gene with a RMI2 domain that may serve the function of ParB.

The plasmids also encode a SMC chromosome segregation ATPase within a 7-gene cluster (ORFs 115-121). This protein can preside over cell-cycle checkpoints [21]. The cluster also encodes another AAA-ATPase common in archaea, together with a DNA primase that structurally aligns very well with the eukaryotic-type DNA primase of *Pyrococcus furiosus* (**Figure 4C**). We infer it synthesizes an RNA primer required for the onset of DNA replication, indicating its potential importance in a vector constructed for genetic engineering. A putative DEAD/DEAH box RNA helicase (ORF34) may remodel RNA structures and RNA–protein complexes.

The plasmids encode a nucleoid protein MC1 (ORF157) that is homologous to eukaryotic histones. The predicted structure aligns well with MC1 from *Methanosarcina thermophila*, but it possesses an additional N-terminal region that is largely unstructured (**Figure 4D**). We conclude that the HMp nucleoid protein MC1 is likewise responsible for plasmid genome compaction while allowing replication, repair and gene expression.

The plasmids appear to encode multiple proteins that could be involved in DNA recombination. One is a helicase with an N-terminal UvrD helicase domain and a C-terminal PD-(D/E)XK nuclease domain (ORF40) that is also found in Cas4 nucleases. This protein was found on several HMp plasmids, as well as the genome of a large plasmid of the methanogen *Methanomethylovorans hollandica* (**Figure S6C**). ORF45 and ORF128 encode HNH-endonucleases that could stimulate recombination. ORF128 has an RRXRR motif, an architecture common to some CRISPR-associated nucleases (COG3513) [22]. Furthermore, the plasmids encode a recombination limiting protein RmuC (ORF176).

The plasmids encode other genes involved in nucleotide processing. This set includes a 5-gene cluster encompassing a putative AAA-ATPase (COG1483, ORF149), two genes of unknown function, a nuclease (ORF144), and a helicase with a similar architecture to the RNA polymerase (RNAP) associated protein RapA. RapA reactivates stalled RNAP through an ATP-driven back-translocation mechanism, thus stimulating RNA synthesis [23]. Furthermore, HMp_v1 encodes a large (1550 AA) protein with an N-terminal ATPase domain and a C-terminal HNH endonuclease domain (ORF39). Interestingly, this latter domain is preceded by a 330 bp region that encodes 11 repeated [PPEDKPPEGK] amino acid sequences that are predicted to introduce intrinsic disorder. The C-terminal region resembles the histone H1-like DNA binding protein and inner and outer membrane linking protein TonB. We speculate that the repeat region facilitates binding of the ATPase/endonuclease to other interaction partners (nucleic acid or protein).

### Transporters and membrane proteins

HMp_v2 encodes 15 membrane proteins and 3 extracellular proteins, whereas the larger HMp_v1 carries 25 predicted membrane proteins and 4 extracellular proteins. This difference is due to one large genetic island in HMp_v1 that encodes several transporters. There is a single gene encoding a Fe^2+^/Mn^2+^ transporter (ORF38) that is also found in some Asgard archaea and bacteria and a region spanning ORFs 54-79 that encodes several transport systems. First, a putative CbiMNQO Co^2+^/Ni^2+^ transporter composed of three membrane subunits and a soluble subunit whose expression could be controlled by the preceding NikR regulator. Second, an amino acid permease (ORF61) whose expression could be regulated by an accompanying HrcA. Third, a second CbiMNQO Co^2+^/Ni^2+^ transporter with similar architecture, whose expression may be controlled by an Ars regulator. Another NikR (ORF67) follows and several proteins with the same DUF3344 that are predicted to be located extracellularly and are likely cell surface proteins (ORF68, ORF70). The genetic region is completed by another ABC-transporter that could be a biotin transporter, since two subunits resemble EcfT and EcfA1/2 (ORFs 72-75). This region appears not to be present in the coexisting *Methanoperedens*, indicating the potential for the plasmids to augment their host’s metabolism.

Two gene clusters flanked by transposases encode two putative membrane complexes. The first includes a secretion ATPase VirB11 and a 7-TMH bearing membrane protein (**Figure S7**). The combination is reminiscent of a system for DNA transfer between *Sulfolobus* cells [24]. The second includes multiple membrane proteins with features suggestive of binding DNA/RNA/proteins and/or lipoproteins and a HerA helicase (ORF112) of unknown localization that possesses a domain found in conjugative transfer proteins. This second cluster could be involved in extrusion of DNA.

The HMp_v1 plasmid encodes four tetratricopeptide repeat proteins (TPR). One is a membrane protein and two are membrane-attached and cytoplasmically orientated TPRs. The fourth is a soluble protein that is accompanied by a small 3-TMH bearing membrane protein and possibly tied to membrane processes in the host. The TPR domains facilitate protein-protein interactions and are for example required for PilQ assembly of the type IV pilus. TPR4 may be involved in the homologous archaeosortase system that cleaves the signal peptide and replaces it with another modification. The HMp_v2 plasmid carries two presumably protein binding pentapeptide repeat-containing proteins (v2 ORFs 16-17).

### Proteins involved in cell protection

Both plasmids encode a dCTP deaminase (ORF156), which preserves chromosomal integrity by reducing the cellular dCTP/dUTP ratio, preventing incorporation of dUTP into DNA. HMp_v1 also encodes a diphtine-ammonia ligase (DAL) (**Figure 4E**) that catalyses the last step of a post-translational modification of the elongation factor eEF2 during ribosomal protein synthesis [25]. A glyoxalase (**Figure 4F**) can convert cytotoxic α-keto aldehydes into nontoxic α-hydroxycarboxylic acids [26]. ArsR may regulate expression of a peptide methionine sulfoxide reductase (MsrA, **Figure 4G**). MsrA repairs oxidative damage to methionine residues arising from reactive oxygen species and reactive nitrogen intermediates [27].

### Expression of plasmid genes

We used metatranscriptomics to determine which of the genes of the high abundance plasmids v2, v5 and the lower abundance v1 are most important to the *Methanoperedens* growing in the bioreactors. Metatranscriptome reads from Bioreactor 1 or 2 were stringently mapped onto all contigs of each respective bioreactor. Read counts were normalized to the gene length and genes were considered expressed that had at least 0.5 mapped reads. We found reads that mapped uniquely onto all three plasmid genomes, and the high coverage plasmids had higher normalized read counts (**Table S2**).

Twenty of the 178 genes of the low coverage plasmid HMp_v1 were expressed. Most highly expressed were the gene encoding the MTH865 protein, which has been structurally characterized but lacks a known function [28], and its accompanying genes (ORFs 30-31). Also highly expressed were the first Co^2+^/Ni^2+^ transporter and two preceding genes. One component of the putative biotin transporter was also expressed and the nucleoid protein MC1. This suggests that this plasmid facilitates or enhances the uptake of Co^2+^/Ni^2+^ and possibly biotin.

Of the 158 genes of the high coverage plasmid HMp_v2 in Bioreactor 2, 103 were expressed. The highest expression of genes (≥100 normalized reads) with functional annotations was observed for ParA and its genetic context, the dCTP deaminase and MTH865. Moderate expression (≥ 10) was observed for genes encoding MsrA and its regulator, as well as the glyoxalase. Thus, we infer that this plasmid is actively conferring protection from oxidative stress and cytotoxic compounds. Genes that only showed low expression are mostly important for plasmid maintenance. Interestingly, the origin of replication proteins of all plasmids were not expressed. However, we detected expression of the OriC adjacent gene, encoding a hypothetical protein which has a P-loop fold. This suggests that this could be an important component in plasmid replication, possibly performing ATP hydrolysis (ORF186).

Of the 164 genes of HMp_v5 from Bioreactor 1, 104 were expressed. The most highly expressed genes of HMp_v5 are the first gene encoding a hypothetical small protein and the last gene encoding the small subunit GroES (Chaperonin Cpn10) of a 3-gene cluster that includes GroEL. This chaperonin system is crucial for accurate protein folding [29]. Other highly expressed genes (≥ 500 normalized reads) encode a HrcA regulator which could enable expression of the equally expressed, adjacently encoded 50S ribosomal protein uL16 and translation initiation factor 2 subunit beta, as well as a rubrerythrin. Furthermore, another ArsR regulator, a putative integrase and a two-component system with resemblance to FleQ, a transcriptional activator involved in regulation of flagellar motility, were highly expressed.

Moderately expressed (≥ 50 normalized reads) were proteins involved in toxin-antitoxin systems, an archaeal translation initiation factor, two adjacently encoded TIR-like nucleotide binding proteins located next to the protein involved in replication initiation and proteins involved in cell growth and apoptosis (IMPDH ParBc_2). Overall, the main function of HMp_v5 may be to ensure protein maturation and regulate DNA processes including transcription and translation.

### Plasmid specificity of proteins

To further understand how the plasmid inventories may augment or overlap with those of the host *Methanoperedens* we performed protein family clustering using a protein dataset composed of 96,548 proteins from the HMp plasmids and *Methanoperedens* chromosomes (**Table S3**). Also included were proteins from Borgs and a small set of reference proteins from plasmids of methanogens. The hierarchical clustering revealed that the plasmid proteomes clustered distinct from *Methanoperedens* and Borg proteomes (**Figure 5A, Figure S8**). Of the 1,079 plasmid proteins, 882 (82%) clustered into 504 subfamilies. The majority of plasmid-encoded proteins had homologues in the *Methanoperedens* genomes (80%). The number of protein subfamilies exclusively shared between plasmids and their host *Methanoperedens* was slightly higher (18%) than for *Methanoperedens* without plasmids (14%) (**Figure 5B, Figure S8**).

**Figure 5.**
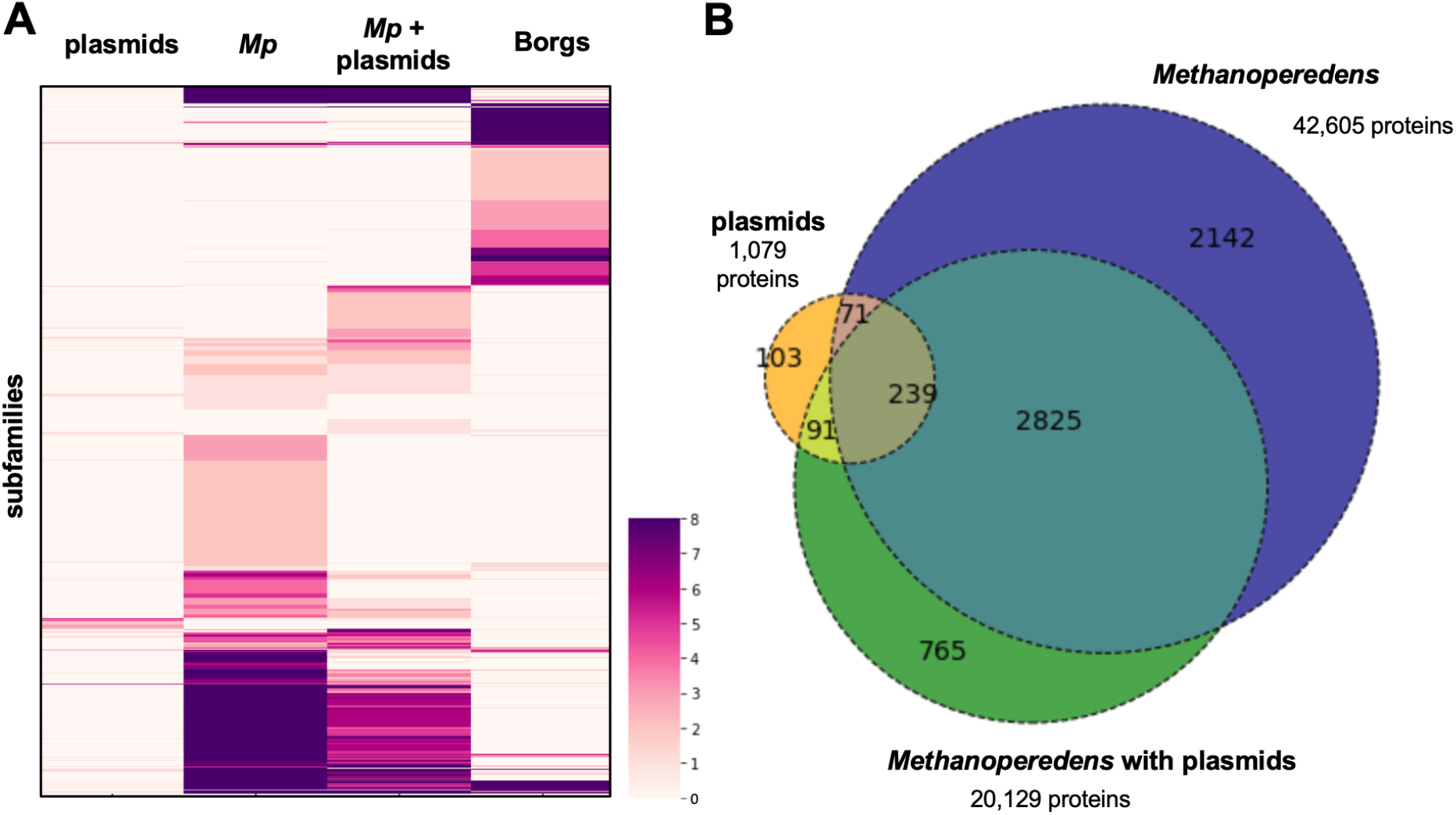
Protein clustering analyses and distribution of protein subfamilies across main elements. **A.** Heatmap showing protein subfamilies (subfamilies ≥ 8 are all shown in dark purple). **B.** Subfamily distribution across plasmids, *Methanoperedens* with plasmids and *Methanoperedens* without plasmids.

Eighteen percent of the plasmid proteins clustered into subfamilies that were unique to the plasmids, and forty-one percent were in subfamilies with only a few non-HMp homologs (**Table 2, Table S3**). Many plasmid-enriched proteins are implicated in DNA replication and repair, including the Cdc24 protein and the DNA primase, and they were actively expressed in HMp_v2. A surprising finding was that there are no homologues on NCBI for a large surface protein that is only found on four HMp versions.

**Table 2.** Protein subfamilies enriched in plasmid proteins. Numbers in brackets indicate the number of subfamily members per plasmid.

| subfamily | PFAM annotation | description | # plasmid members | # total members | % enr. | plasmids | v1 ORF |
| --- | --- | --- | --- | --- | --- | --- | --- |
| subfam0052 | Exo_endo_phos | nuclease | 3 | 3 | 100 | v1, v2, v5 | 144 |
| subfam0053 | DUF87 AAA_10 | HerA helicase | 3 | 3 | 100 | v1, v2, v8 | 112 |
| subfam0085 | FYDLN_acid |  | 2 | 2 | 100 | v1, v2, v8 | 177 |
| subfam2357 | AAA_33 |  | 2 | 2 | 100 | v1, v2 | 186 |
| subfam2361 | Cas_Cas02710 | CRISPR-associated protein | 2 | 2 | 100 | v5, v8 |  |
| subfam2619 | AAA_26 | ParA2 | 3 | 3 | 100 | v1, v2, v8 | 102 |
| subfam3152 | Big_3_3 | surface protein | 4 | 4 | 100 | v1, v2, v4, v8 | 25 |
| subfam6300 | CDC24_OB3 | Cdc24 | 3 | 3 | 100 | v1, v2, v8 | 94 |
| subfam7697 | LexA_DNA_bind | LexR | 3 | 3 | 100 | v1, v2, v8 | 22 |
| subfam7758 | KdpD DUF2478 | secretion ATPase VirB | 4 | 4 | 100 | v1, v2, v4, v8 | 89 |
| subfam0185 | RuvC_1 | transposase | 3 | 13 | 23 | v1 (2), v3 (5), v5 | 43, 88 |
| subfam0188 | DNA_primase_S | DNA primase | 3 | 4 | 75 | v1, v2, v8 | 120 |
| subfam0434 | DUF1156 MethyltransfD12 N6_N4_Mtase |  | 4 | 7 | 57 | v1, v2, v4, v7 | 146 |
| subfam1990 | ThiF | ThiF | 6 | 8 | 75 | v1, v2, v3, v5, v8 (2) | 7 |
| subfam3527 | AAA_11 PhoH |  | 7 | 22 | 32 | v1, v2, v5 (2), v6, v7 (2) | 40 |
| subfam4932 | AAA_11 AAA_12 | UvrD helicase | 5 | 7 | 71 | v1 (2), v2, v3, v8 | 20, 123 |
| subfam5002 | RMI2 | RepA2 | 8 | 65 | 12 | v1 (2), v2 (2), v3, v5, v7, v8 | 3, 169 |
| subfam6245 | YpsA | YpsA | 5 | 7 | 71 | v1, v2, v5, v7, v8 | 18 |
| subfam6302 | DUF3584 Macoillin Cast MAD MAD AAA_13 | chromosome-segregation ATPase | 3 | 6 | 50 | v1, v2, v8 | 117 |
| subfam6760 | NARP1 SPO22 | TPR3 | 3 | 9 | 33 | v1, v2, v3 | 178 |
| subfam6769 | CDC24_OB3 | RepA1 | 6 | 58 | 10 | v1, v2, v3, v5, v7, v8 | 2 |
| subfam7198 | UTP25 DUF1998 | RNA helicase | 4 | 6 | 67 | v1, v2, v4, v5 | 34 |
| subfam7280 | Integrin_beta | Integrin | 4 | 6 | 67 | v1, v2, v3, v8 | 129 |
| subfam7300 | RRXRR RE_Alw26IDE | HNH endonuclease | 6 | 31 | 19 | v1, v7, v8 (4) | 128 |
| subfam7324 | RadC | RadC | 5 | 41 | 12 | v1, v3, v5, v7, v8 | 11 |
| subfam7760 | ChAPs | TPR1 | 4 | 5 | 80 | v1, v2, v5, v6 | 28 |

There were a few instances of plasmid proteins that are auxiliary/linked to central metabolic functions, for example a protein responsible for removing ammonia from glutamine (HMp_v5), a putative cobalamin-independent methionine synthase implicated in AA metabolism (HMp_v7), and a multiheme cytochrome (MHC) that may be important for electron transfer to the final electron acceptor of CH_4_ oxidation (HMp_v7). Other proteins shared with *Methanoperedens* are potentially involved in sensing and signalling. For example, a TPR protein (HMp_v1, v2, v8), a putative nitroreductase which could function in FMN storage (HMp_v8), a phosphoglucomutase/phosphomannomutase possibly tied to glycosylation (HMp_v5), a methyltransferase involved in RNA capping in eukaryotes (HMp_v2, v8), an rRNA methylase (HMp_v5) that could be implicated in post-transcriptional modification, a translation initiation factor (HMp_v5) [30], a peptidyl-tRNA hydrolase involved in releasing tRNAs during translation (HMp_v7) and a phosphoribosyltransferase implicated in stress response (HMp_v7) [31].

HMp_v8 carries two IS200-like transposases and a homologue is also found on the *Methanosarcina barkeri* 227 plasmid (WP_048116267.1). There are two subfamilies that are phage integrases, one of which is also found on *Methanococcus maripaludis* C5 plasmid pMMC501 (WP_010890222.1) and *Methanosarcina acetivorans* plasmid pC2A (WP_010891114.1).

## Discussion

In two laboratory-scale bioreactors and three different natural ecosystems we discovered large, circular plasmids of *Methanoperedens.* To our knowledge, these are the first reported plasmid sequences in the archaeal family *Candidatus* Methanoperedenaceae and the first in ANME archaea. Notably, the deduced hosts for the plasmids are a distinct *Methanoperedens* species group that includes all strains that are currently in laboratory cultures (to our knowledge). For example, “*Ca.* Methanoperedens BLZ2” (Bioreactor 1) carries HMp_v5 and “*Ca.* M. nitroreducens Vercelli” (Bioreactor 2) carries HMp_v2. The plasmids are large compared to most plasmids of methanogens (4,440-58,407 bp). The only exception is the report of a 285,109 bp, plasmid of the obligately methylotrophic methanogen *M. hollandica* DSM 15978 [32].

Although our data indicate that some *Methanoperedens* may carry more than one plasmid, the most abundant (main) plasmid appears to be maintained at an ~1:1 ratio with the host chromosome. Thus, there seems to be coordination of replication of the plasmid and the main chromosome, as has been observed for other archaeal chromosomes and their megaplasmids *[15]*. Maintaining a large plasmid at the same abundance as the main chromosome comes at an energetic cost, suggesting that the plasmids confer cellular fitness or are possibly even essential for the host’s survival.

Interestingly, we identified Orc1/Cdc6 near the origin of replication, but these genes were not expressed at the time of sampling. This could simply be due to the very slow growth rate of *Methanoperedens* in the bioreactors, and concomitantly low replication rates of its plasmids. Since the plasmids do not encode recombinase RadA, this excludes the possibility of origin-less replication initiation via homologous recombination as described for some viruses and archaea [33]. It would however also be possible that the host Orc1/Cdc6 is used to couple replication of the plasmid to chromosome copy number.

There are different versions of the plasmids, but they share elements of a core machinery likely essential for plasmid replication and maintaining DNA integrity. There are also unique functions for different plasmid versions. The observation that HMp_v1 encodes highly expressed genes for Ni^2+^/Co^2+^ transporters is interesting because Ni^2+^ is required for Mcr, the enzyme complex central to methane oxidation, and for the carbon monoxide dehydrogenase/acetyl-CoA synthase. Co^2+^ is part of a complex organometallic cofactor B12, which is essential for the function of methyltransferases [34]. HMp_v2 lacks the genomic island rich in transporters and the metatranscriptomic data indicates that one of its functions is to protect the host from oxidative stress and cytotoxic compounds. HMp_v5 on the other hand predominantly expressed genes tied to protein maturation and regulation of cellular functions, often connected to nucleotide mechanisms. Interestingly, HMp_v5, but not its host’s chromosome, carries the 50S ribosomal protein uL16 and an adjacent gene encoding translation initiation factor 2 subunit β, essential genes for construction of functional ribosomes and translation [35,36]. This suggests that *Methanoperedens* is dependent on the HMp_v5 plasmid, ensuring plasmid retention. The relocation of uL16 to an extrachromosomal element is reminiscent of the relationship in eukaryotes between mitochondrial DNA and nuclear DNA, where many mitoribosomal proteins are encoded in the nuclear DNA. In the case of the plasmid, this control of uL16 could ensure that increased host ribosome production leads to increased translation of plasmid genes.

Based on the phylogenetic analysis, we inferred that plasmids occur in *Methanoperedens* species that do not host Borgs. It was suggested that Borgs are not obviously plasmids [12], but a limitation on their classification was the lack of archaeal plasmids generally, and *Methanoperedens* plasmids, specifically, to compare them to. Here, we find that, in contrast to Borgs, the plasmids do not, or only very rarely (e.g., one MHC), carry genes with a protein function associated with the central metabolism of their host (anaerobic methane oxidation). The observations presented here underline the distinction between Borg extrachromosomal elements and plasmids of *Methanoperedens*.

Although archaeal methanotrophs of the genus *Methanoperedens* have been studied using cultivation-independent [37] and enrichment-based methods [3], many questions regarding their physiology remain. We hope that this discovery of naturally occurring plasmids associated with *Methanoperedens* in stable enrichment cultures, paired with the possibility of editing the genomes of specific organisms in microbial communities [38], is a first step towards developing genetic modification approaches to better understand anaerobic oxidation of methane and potentially to harness this process for agricultural and climate engineering.

## Methods

### Identification of ECEs associated with *Methanoperedens* and manual genome curation

Metagenomic datasets on ggKbase (ggkbase.berkeley.edu) were searched for contigs with a dominant taxonomic profile matching *Methanoperedens*(Archaea; Euryarchaeaota; Methanomicrobia; Methanosarcinales; Candidatus Methanoperedens; Candidatus Methanoperedens nitroreducens). Manual genome binning was performed based on coverage, GC content and contig taxonomy. Plasmids were identified based on marker proteins (Orc1/Cdc6) and whole genome alignments using the progressive Mauve algorithm. Additional plasmids in environmental metagenome datasets were identified by BLAST and verified by genome alignment to a bioreactor plasmid [39]. Manual curation of two plasmid sequences to completion was performed in Geneious Prime 2021.2.2 (https://www.geneious.com). Curation involved piecing together and extending contigs with approximately the same GC content, depth of sampling (coverage), and phylogenetic profile. Sequence accuracy and local assembly error correction made use of read information, following methods detailed in [40]. The final, extended sequences contained identical regions at the termini, and were thus circularized. The start positions of the genomes were chosen based on cumulative GC skew information.

### Nucleic acid extractions from the *Methanoperedens* enrichment cultures and plasmid isolation

DNA and RNA samples were taken from Bioreactor 1 in April 2021. DNA samples were taken from Bioreactor 2 in April 2021 and RNA samples were taken from a subculture of Bioreactor 2 in October 2021. DNA was isolated following the Powersoil DNeasy kit protocol, with the addition of a 10 min bead beating step at 50 s^−1^ (Qiagen, Hilden, Germany). RNA was isolated following the Ribopure-Bacteria kit protocol (Thermo Fisher Scientific, Waltham, US), with the addition of a step homogenizing the cells and a 15 min bead beating step at 50 s^−1^. The metatranscriptomic datasets were constructed from technical replicates (n=4 for Bioreactor 1, n=3 for Bioreactor 2). Plasmids were targeted in a second bioreactor sampling experiment (n=2 for both bioreactors) for which the Plasmid Miniprep kit was used according to manufacturer’s instruction (Thermo Fisher Scientific, Waltham, US), with the addition of a step homogenizing the cells before processing. The metagenomic datasets were constructed from biological replicates (n=2-4).

### Metagenomic and metatranscriptomic assemblies

DNA was submitted for Illumina sequencing at Macrogen or at the in-house facility of Radboud University to generate 150 or 250 bp paired end (PE) reads for metagenomes, and 100 bp PE for metatranscriptomes. Sequencing adapters, PhiX and other Illumina trace contaminants were removed with BBTools and sequence trimming was performed with Sickle. The filtered reads were assembled with IDBA-UD [41] (v1.1.3), ORFs were predicted with Prodigal [42] (v2.6.3) and functionally annotated by comparison to KEGG, UniRef100 and UniProt using USEARCH [43] (v10.0.240). Metagenomic and metatranscriptomic reads were assembled with IDBA-UD.

The metatranscriptomic reads of Bioreactor 1 or 2 and replicate 1 or 2 were mapped against the all contigs from the same sample using BBMap and a stringent mapping where reads had to be 99% identical to map (minid=0.99 ambiguous=random). The mapped reads per gene were calculated with featureCounts (--fracOverlapFeature 0.1). The resulting read counts were normalized to gene length and are given as the number of reads per 1,000 bp. Normalized reads ≥ 0.5 were considered expressed.

### Nucleotide alignments and phylogenetic tree construction

Whole genome alignments were done in Geneious using the progressiveMauve algorithm when aligning complete genomes or single contigs, or MCM algorithm when aligning genomes on multiple contigs. RpS3, UvrD, RadC and helicase/nuclease genes were aligned with MAFFT (v7.453), trimmed with trimal (-gt 0.2) [45] (v1.4.rev15) and a maximum-likelihood tree was calculated in IQ-Tree (-m TEST -st AA -bb 1000 -nt AUTO -ntmax 20 -pre) [46]. The trees were visualized and decorated in iTOL [47]. tRNA alignments were constructed by adding predicted host and phage tRNAs to archaeal tRNA alignments from GtRNAdb release 19 [48] with the add option of mafft [44] (v7.453). Using these alignments tRNA phylogenies were constructed with IQ-tree, using the automatic model finder and 1000 bootstrap replications [46]. The trees were visualized and decorated in iTOL [47].

### Structural, functional and localization predictions

Proteins were profiled using InterProScan [49] (v5.51-85.0) and HMMER (https://hmmer.org) (v3.3, hmmsearch) against the PFAM (--cut_nc) and KOFAM (--cut_nc) HMM databases [50,51]. TMHs were predicted with TMHMM [52] (v2.0) and cellular localization using PSORT [53] (v2.0, archaeal mode). tRNAs were searched with tRNAscan [48] (v.2.0.9) and rRNAs with SSU-ALIGN [54] (v0.1.1). Plasmid protein structures were modelled using AlphaFold2 [55] via a LocalColabFold [56,57] (--use_ptm --use_turbo --num_relax Top5 --max_recycle 3), visualized and superimposed onto PDB structures using PyMOL [58] (v2.3.4). Structure-based homology search was performed in PDBeFold [59]. Plasmid comparison figure was generated with clinker [60] (v0.0.21).

### Protein family clustering

A dataset of 96,548 proteins was constructed using the elements in the project folders listed in the Data availability statement. These cover all 8 HMp versions, 4 complete Borg genomes, additional incomplete Borg genomes, and *Methanoperedens* genomes. This core dataset was supplemented with reference genomes comprising protein sequences from plasmids of methanogens, and from “*Ca*. Methanoperedens nitroreducens”, “*Ca*. M. ferrireducens”, “*Ca*. M. manganicus” and “*Ca*. M. manganireducens” (Table S4). All proteins were clustered into protein subfamilies using MMseqs [61] and HMMs were constructed from these subfamilies using HHblits [62] as previously described [63]. They were then profiled against the PFAM database by HMM-HMM comparison using HHsearch [64] and protein subfamilies enriched in plasmid proteins were determined as described previously [65].

### Replication prediction by GC skew analysis

GC skew and cumulative GC skew were calculated as described previously [66].

## Acknowledgements

Funding for this research was provided by a DFG fellowship for MCS (Project Number: 447383558), the Soehngen Institute of Anaerobic Microbiology Gravitation program through grant 024.002.002 by the Dutch Science Foundation and the Innovative Genomics Institute at UC Berkeley. The Ministry of Economy, Trade and Industry of Japan funded a part of the work as “The project for validating near-field system assessment methodology in geological disposal system” (2020 FY, Grant Number: JPJ007597). Shufei Lei and Jordan Hoff for bioinformatics support and Justin Smith, Luis Valentin Alvarado, Susan Mullen, Kenneth Williams, Karthik Anantharaman and Basem Al-Shayeb for their contributions to field work and generation of sequence datasets.

## Author contributions

The study was designed and performed by MCS, JFB, HO and CW. HO and CW established, maintained and sampled the bioreactors. HO extracted the DNA and RNA and obtained sequence datasets. JWR and JFB provided the Corona Mine dataset and YA provided the Horonobe metagenomic dataset. Genome, proteome, phylogenetic and transcriptome analyses were performed by MCS. JWR assisted with computational analyses. RS contributed to the data handling and supported the bioinformatic analyses. JFB performed the binning and carried out the manual genome curation. PP contributed to the protein functional analysis. MCS and JFB wrote the manuscript with input from all authors.

## Data availability

Newly released sequences used in this manuscript are available via: https://ggkbase.berkeley.edu/P_Mp/organisms. The publicly available dataset used for other *Methanoperedens* and Borg analyses can be accessed via: https://ggkbase.berkeley.edu/BMp/organisms.

## Supplementary Data

**Figure S1.**
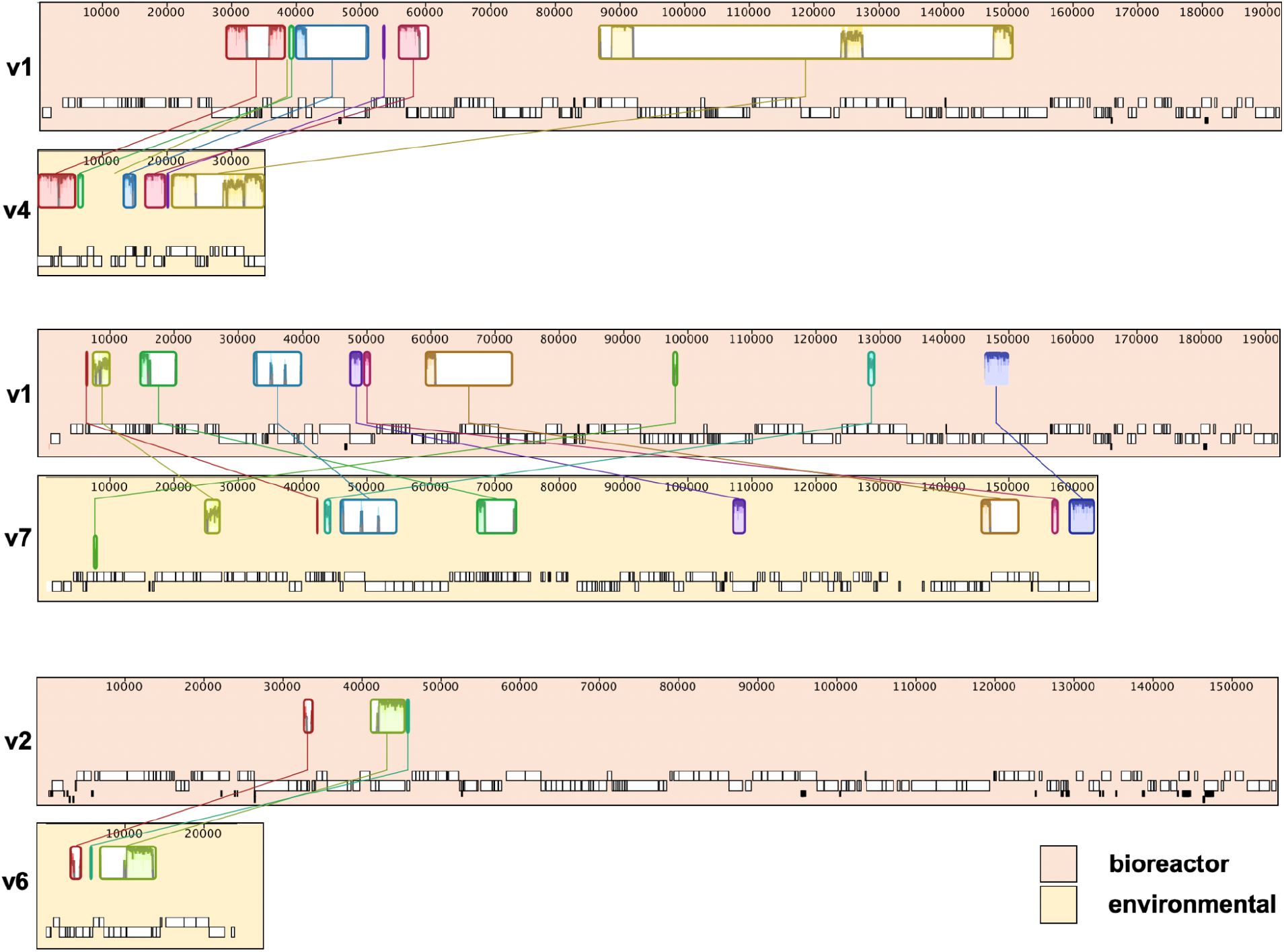
Nucleotide alignment of plasmid fragments from environmental samples to the curated v1 or v2 plasmid genome.

**Figure S2.**
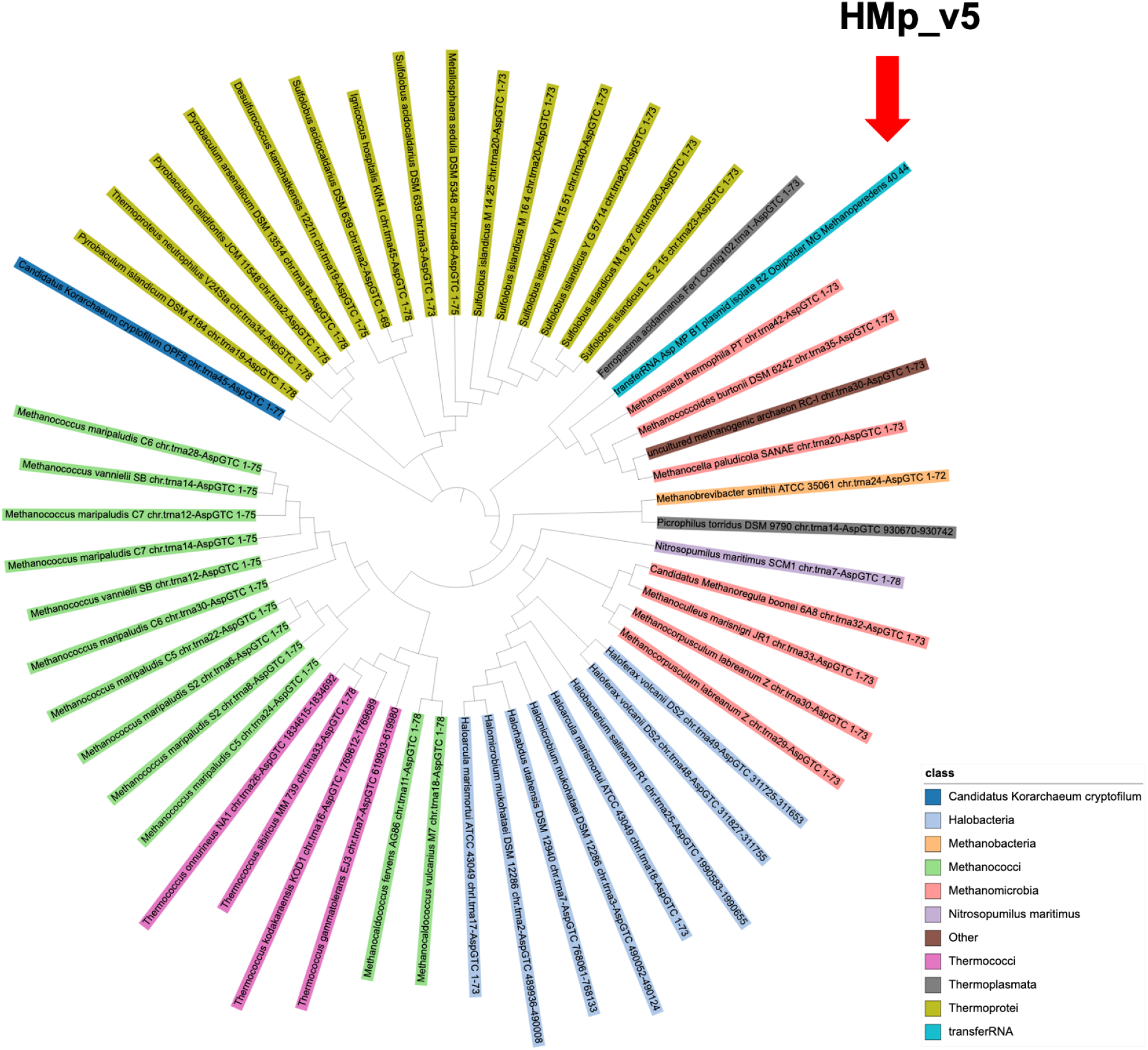
Phylogenetic tree of archaeal Asp tRNAs. Asp tRNA of HMp_v5 falls within a clade of other Asp tRNA from *Methanomicrobia.* It is absent from the host genome.

**Figure S3.**
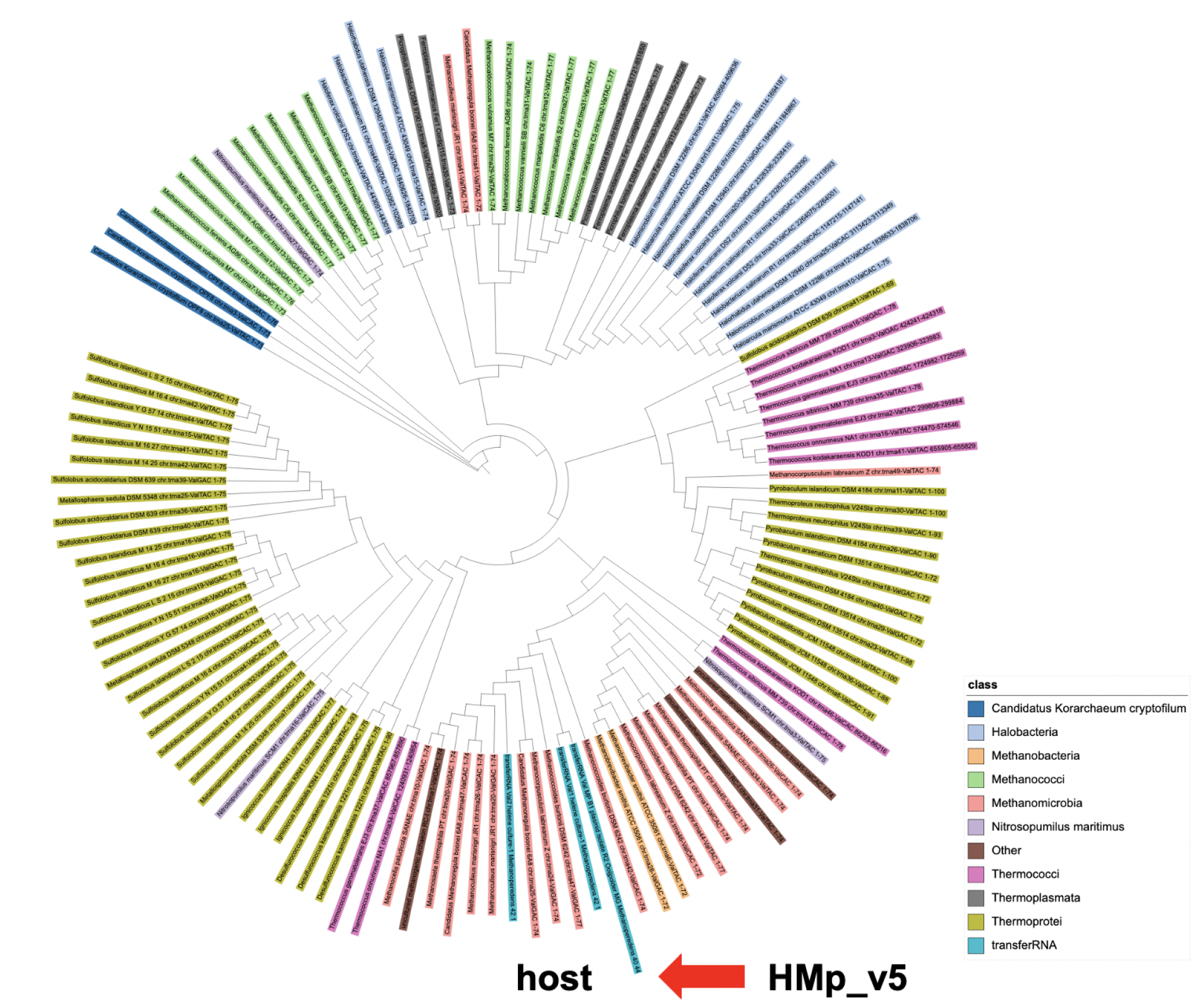
Phylogenetic tree of archaeal Val tRNAs. Val tRNA of HMp_v5 falls within a clade of other Val tRNA from *Methanomicrobia,* including the two from the host *Methanoperedens*.

**Figure S4.**
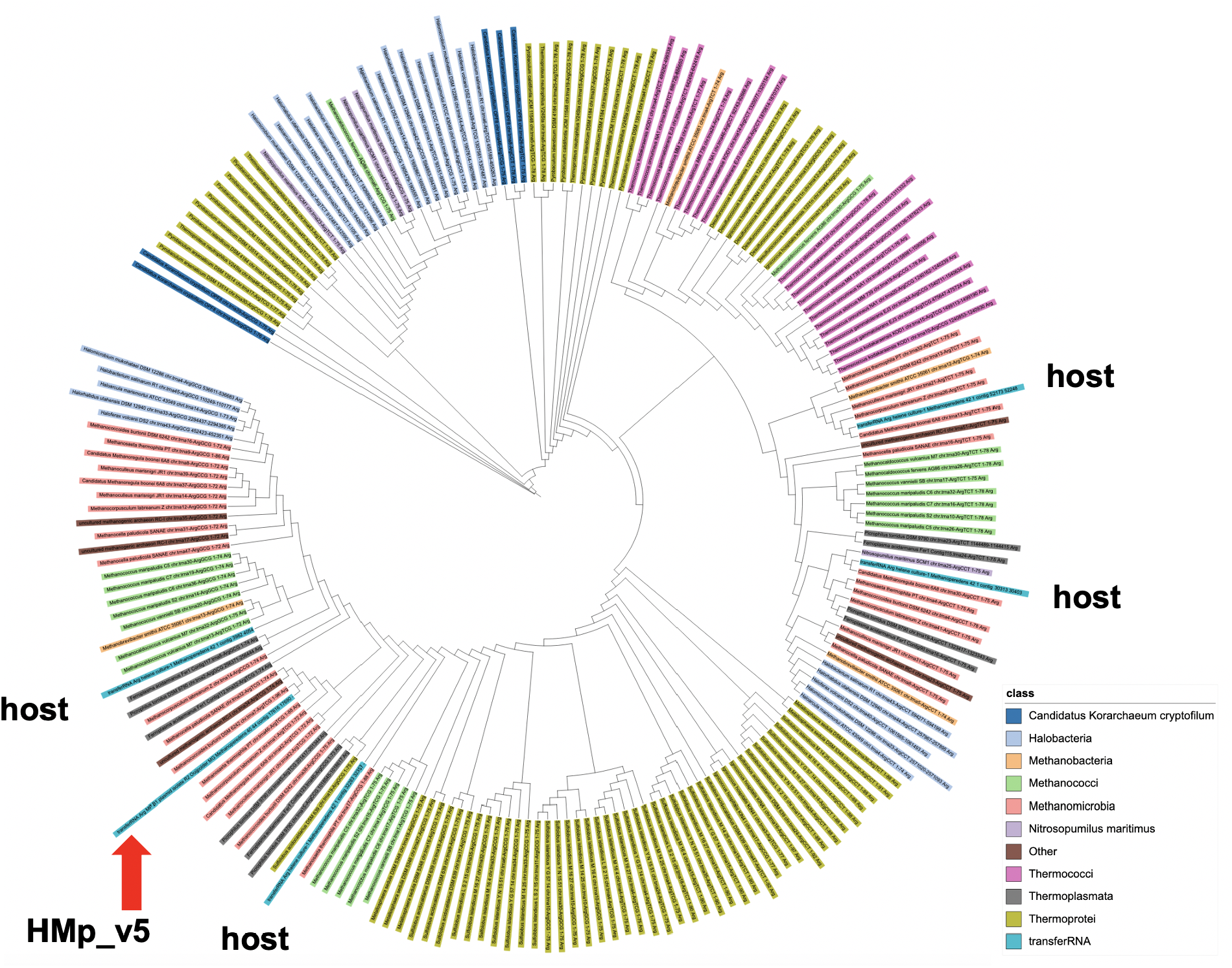
Phylogenetic tree of archaeal Arg tRNAs. Arg tRNA of HMp_v5 falls within a clade of other Asp tRNA from *Methanomicrobia.* The host has four Arg tRNA, but with different anticodon types.

**Figure S5.**
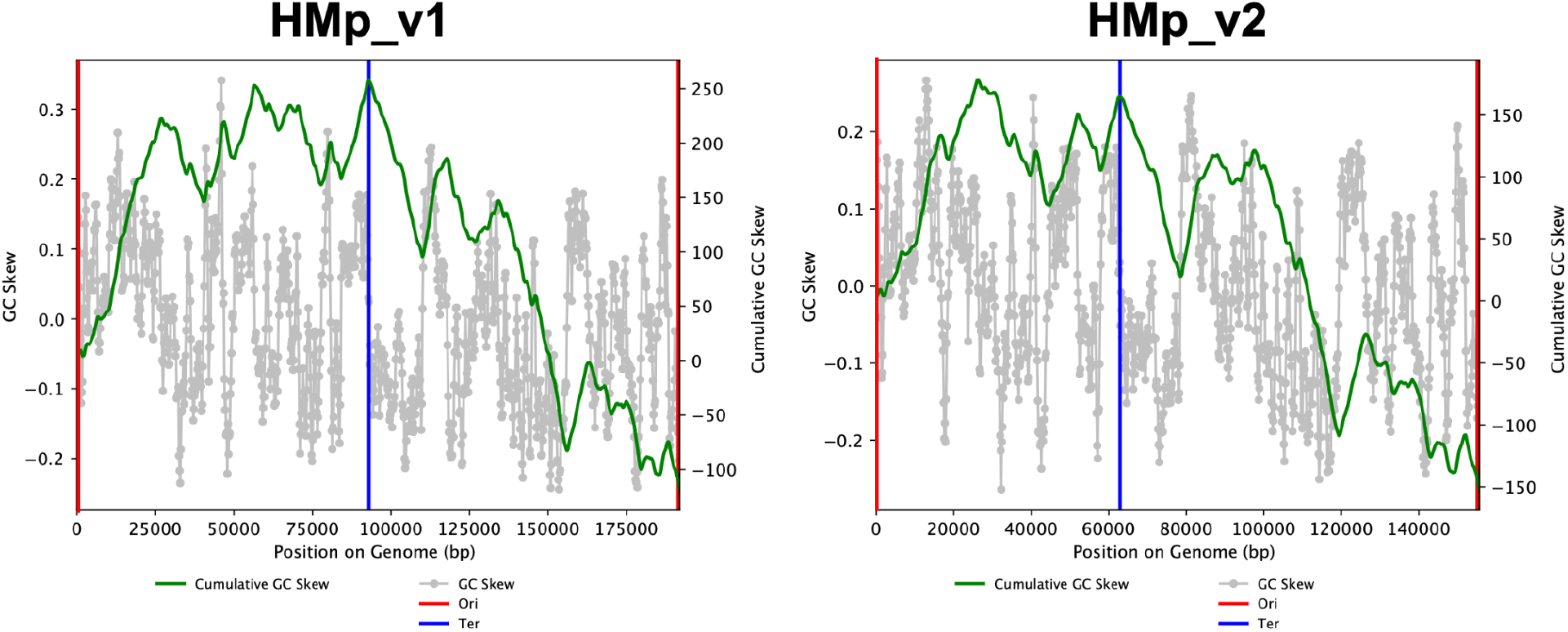
GC skew analysis of HMp_v1 and HMp_v2. Origin (Ori) of replication and termini (Ter) were calculated from cumulative GC skew.

**Figure S6.**
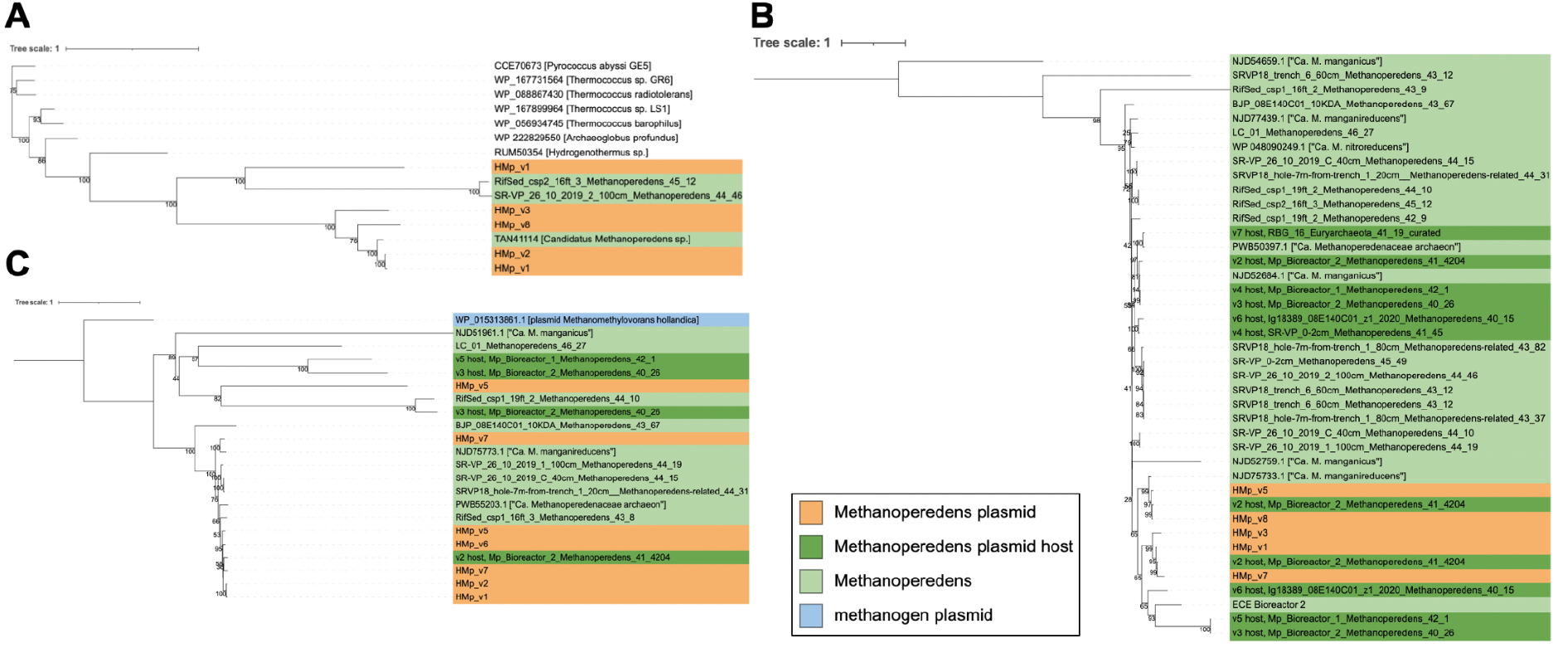
Phylogenetic tree of plasmid enriched protein subfamilies comprising UvrD helicase, RadC and a helicase/nuclease. **A.** Unrooted tree of UvrD helicase proteins (subfamily4932+selected top Blastp hits). Plasmid UvrD cluster together, are absent in the host genomes, but cluster with *Methanoperedens* genomes **B.** Phylogenetic tree of RadC proteins (subfamily7324) rooted on NJD54659.1 from “*Ca.* M. manganicus”. Plasmid proteins cluster together with a group of homologues from plasmid host *Methanoperedens* and an unclassified extrachromosomal element from Bioreactor 2. **C.** Phylogenetic tree of helicase/nuclease protein (subfamily3527) rooted on protein from *M. hollandica* plasmid. The plasmid proteins cluster together. The second versions on v5 and v7 are more related to proteins found on *Methanoperedens* genomes.

**Figure S7.**
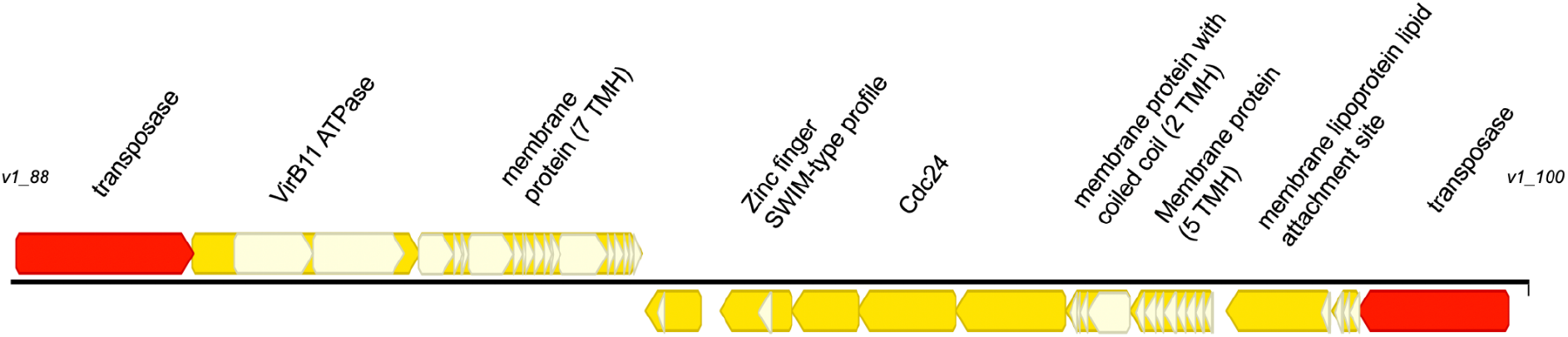
Two gene clusters on HMp_v1 encoding putative membrane complexes.

**Figure S8.**
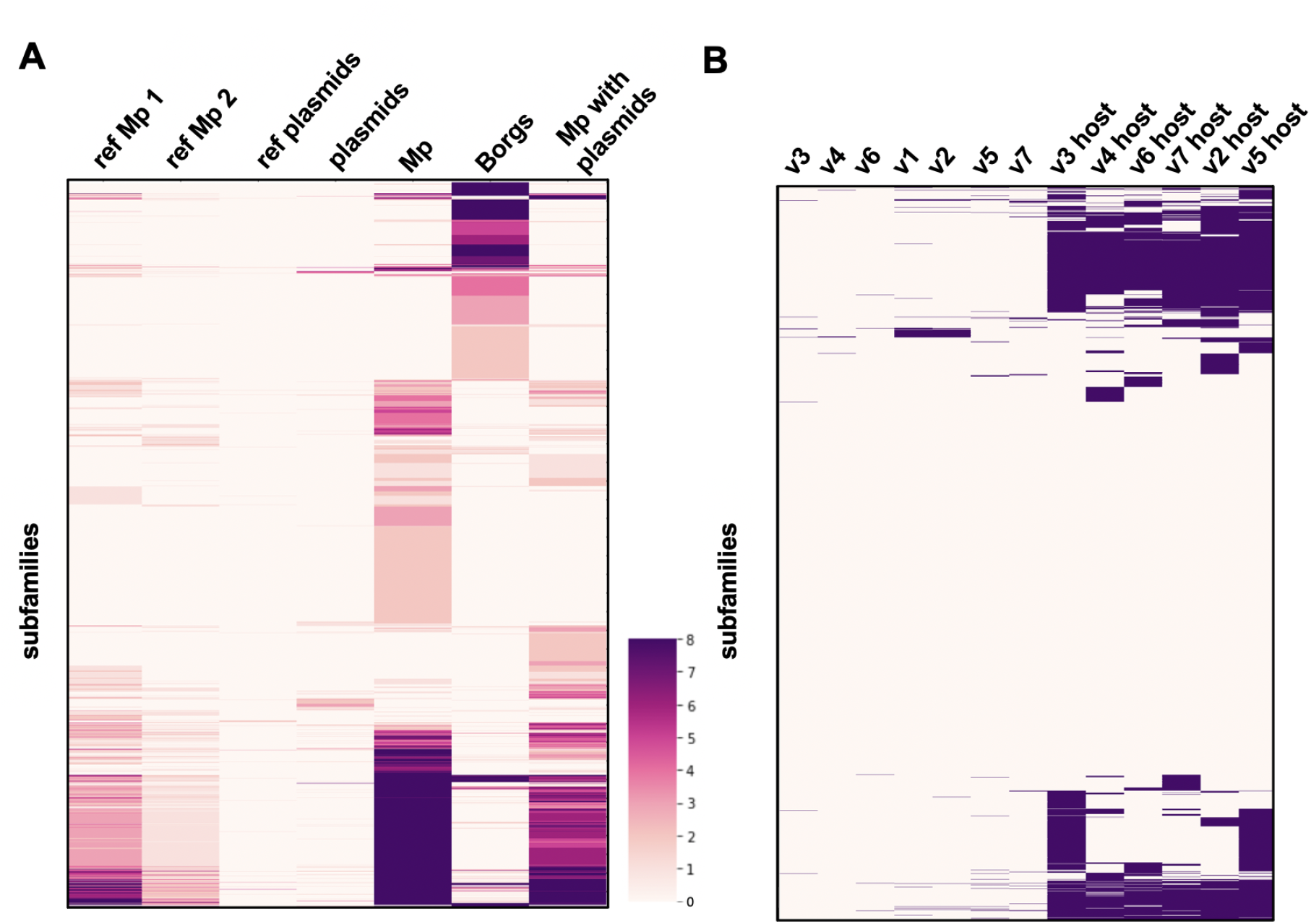
Protein subfamilies of plasmids, *Methanoperedens* with or without plasmids, Borgs, reference *Methanoperedens* and reference plasmids. **A.** Heatmap showing protein subfamilies (subfamilies ≥ 8 are all shown in dark purple). Reference *Methanoperedens* group 1 (ref Mp 1) comprise proteomes of “*Ca*. M. nitroreducens”, “*Ca*. M. ferrireducens”, “*Ca*. manganicus”, ref Mp 2 is proteome of “*Ca*. M. manganireducens”, reference plasmids comprise 8 plasmid proteomes of methanogens. **B.** Presence-absence map of plasmid proteomes (v1-v7) and their inferred *Methanoperedens* host’s proteomes.

## Notes

https://ggkbase.berkeley.edu/P_Mp/organisms

https://ggkbase.berkeley.edu/BMp/organisms

### Competing Interest Statement

J.F.B. is a founder of Metagenomi.

